# Vitamin C Binds to SARS Coronavirus-2 Main Protease Essential for Viral Replication

**DOI:** 10.1101/2021.05.02.442358

**Authors:** Tek Narsingh Malla, Suraj Pandey, Luis Aldama, Dennisse Feliz, Moraima Noda, Ishwor Poudyal, George N. Phillips, Emina A. Stojković, Marius Schmidt

## Abstract

There is an urgent need for anti-viral agents that treat and/or prevent Covid-19 caused by SARS-Coronavirus (CoV-2) infections. The replication of the SARS CoV-2 is dependent on the activity of two cysteine proteases, a papain-like protease, PL-pro, and the 3C-like protease known as main protease Mpro or 3CLpro. The shortest and the safest path to clinical use is the repurposing of drugs with binding affinity to PLpro or 3CLpro that have an established safety profile in humans. Several studies have reported crystal structures of SARS-CoV-2 main protease in complex with FDA approved drugs such as those used in treatment of hepatitis C. Here, we report the crystal structure of 3CLpro in complex Vitamin C (L-ascorbate) bound to the protein’s active site at 2.5 Ångstrom resolution. The crystal structure of the Vitamin C 3CLpro complex may aid future studies on the effect of Vitamin C not only on the coronavirus main protease but on related proteases of other infectious viruses.

---

The Covid-19 pandemic, caused by a novel severe acute respiratory syndrome (SARS) coronavirus 2, has paralyzed public life globally. It has resulted in excess of 500,000 deaths in the US alone as of April 2021. Although potent vaccines are now available, they are optional, not tested in children and potentially not as effective against new viral mutant variants emerging world-wide (LITER). The need for an effective inexpensive treatment and prevention of Covid-19 still exists as large number of infections with potentially severe or lethal outcomes are still reported daily. SARS CoV-2 is a large, enveloped single-stranded RNA betacoronavirus (Cui et al., 2019). The viral RNA encodes two open reading frames that, generates two polyproteins pp1a and pp1ab8. The polyproteins are processed by two viral cysteine proteases: a papain-like protease (PLpro) and a 3C-like protease, also referred to as the main protease (Mpro) or 3CLpro, that cleave multiple sites to release non-structural proteins (nsp1-16). These non-structural proteins form the replicase complex responsible for replication and transcription of the viral genome and have led to 3CLpro and PLpro being the primary targets for antiviral drug development. The 3CLpro is a chymotrypsin-like cysteine protease with a Cys/His dyad in the active site (Fig. 1a). The 3C refers to region 3C in the genome of the *picornaviridae* family, where a similar protease is found (Ramajayam et al., 2011). In coronaviruses, the 3CLpro is located in the non-structural protein (NSP) 5 coding region of the virus RNA genome (Wu et al., 2020).

**Figure 1.**
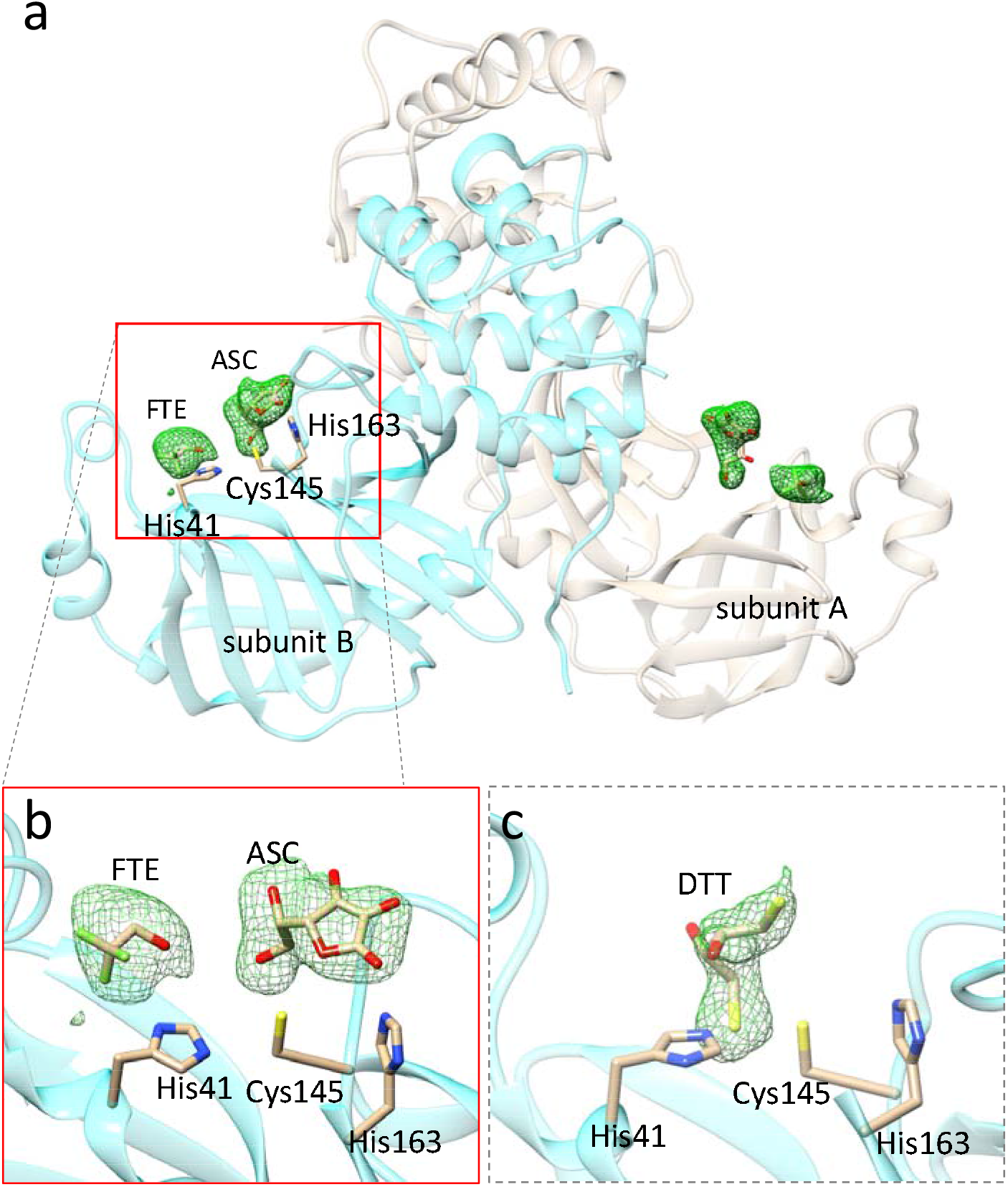
Vitamin C and DTT bound in the active site of SARS CoV-2 3CLpro. (a) The 3CLpro dimer in the asymmetric unit of the orthorhombic crystals soaked with L-ascorbate. Polder difference electron density (Liebschner et al., 2017) in the active sites is shown in green (contour: 2.5 sigma). (b) The L-ascorbate (ASC) in subunit B. The ASC interacts with the catalytic Cys-145 and is stabilized by a hydrogen bond to His-163. A trifluoroethanol (FTE) is located close to the ASC. (c) Dithiothreitol (DTT) is observed in monoclinic (C2) crystal form. It does not bind to Cys-41. It rather binds to His-41 forming a sulfenamide. Difference electron density maps shown in (a) - (c) are obtained after refining the 3CLpro without the addition of ligand. The ascorbate and DTT explain the additional electron density.

High throughput structure based drug discovery experiments on 3CL-pro were recently conducted by large groups of scientists at synchrotrons such as Petra III (Gunther et al., 2021), Diamond (Douangamath et al., 2020) and NSLS-II (to be published) among others. This approach was also rapidly deployed after the 2002 SARS-CoV-1 outbreak, with earlier work by the Hilgenfeld group on Mpro (3CLpro) of coronarviruses leading to crystal structures of SARS-CoV-2 Mpro and inhibitor complexes (Zhang, Lin, Kusov, et al., 2020; Zhang, Lin, Sun, et al., 2020). Active sites of coronavirus Mpro are well conserved and those of enteroviruses (3Cpro) are functionally similar: thus, providing an excellent opportunity to develop broad-spectrum antivirals with structural biology approach. However, the most inhibitors investigated so far are marginally water soluble. They have to be added to the crystals in an organic solvent, or they can be co-crystallized. Their relative scarcity, potential toxicity and unspecificity, and a potentially expensive price tag prevents their wide-spread use. Inexpensive, water soluble, and readily available drugs to combat Covid-19 are urgently required.

Vitamin C has been shown to have antiviral activity for more than half a century, including work of the two-times Nobel Laureate Linus Pauling (Pauling, 1970). Vitamin C is a six-carbon compound (Scheme 1) and a potent antioxidant. It is available in synthetic form and found naturally in Indian gooseberry, citrus fruits and green leafy vegetables. The revival of interest in Vitamin C therapy for acute inflammatory disorders, grounded in sound biological rationale, follows decades of research. Emerging literature suggests that Vitamin C may also play an adjunctive role in the treatment of a variety of viral infections (Colunga Biancatelli et al., 2020; Fowler Iii et al., 2017). A number of observers including Linus Pauling have suggested in the past that Vitamin C in high dosages is directly virucidal (Furuya et al., 2008; Harakeh et al., 1990; Klein, 1945; Pauling, 1970). This assumption was based on in vitro studies, where very high doses of Vitamin C, in the presence of free copper and/or iron, has virucidal activity, presumably through the generation of hydrogen peroxide and other radical species (Furuya et al., 2008; Klein, 1945).

**Scheme 1.**
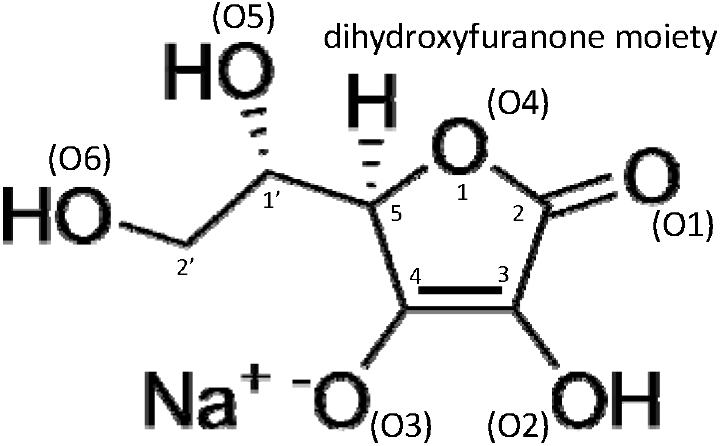
The chemical structure of sodium L-ascorbate, sodium 5-[(1S)-1,2-dihydroxyethyl]-3-hydroxy-4-oxo-furan-olate. The crystallographic annotation of atoms is shown in brackets for oxygens that can be involved in hydrogen bonds.

A recent clinical study of injecting high doses of vitamin-C (> 20 g) intravenously to combat Covid-19 yielded promising results (Zhao et al., 2021). In another study 8 g of Vitamin C was administered orally over three meals together with zinc (Thomas et al., 2021). The results were inconclusive although a positive trend is apparent. So far, it could not be demonstrated that Vitamin C is directly virucidal against Covid-19 *in vivo*.

In order to investigate the binding of Vitamin-C directly to the CoV-2 3CLpro, we co-crystallized CoV-2 3CLpro with low concentrations (5 mmol/L) of L-ascorbate and soaked these crystals before freezing for 3 min in 120 mmol/L ascorbate (Supplementary Material). Difference electron density that can be assigned to ascorbate appears in the active sites near Cys-145 and His-163 of both subunits (Fig. 1). The L-ascorbates only weakly interact with Cys-145 (between 3 Å and 3.7 Å to the sulfur, Tab. 1) but form a tight hydrogen bonding network with Asn-142, Gly-143(N), His-163 and Gln-166 (Fig. S1 and Tab. 1). In addition to the L-ascorbate electron density, an additional density feature is present in the active site. The pillow-like electron density (Fig. 1 b) is interpreted with a trifluoroethanol (FTE) that has been provided as an additive to the crystallization buffer. The interaction of the FTE with the nearby L-ascorbate is weak as the closest distance is larger than 4 Å (Tab. 1). Fig. 2 compares the L-ascorbate position to the position of compound 13b in the active site of the 3CLpro. Compound 13b is a potent inhibitor of the 3CLpro (Zhang, Lin, Kusov, et al., 2020). The positions of the two compound 13b oxygens O40 and O41 in the 3CLpro are close to the C2’ oxygen of the ascorbate (Fig. 2 a). The dihydroxyfuranone ring (Scheme 1) of the ascorbate occupies a similar position as the moiety P1 of compound 13b (Fig. 2 a). The FTE oxygen position is reflected by a structural water found in the compound 13b complexed 3CLpro structure. 3CLpro can also be crystallized in the presence of dithiotreitol (DTT) (Zhang, Lin, Sun, et al., 2020). DTT binds with one of its sulfurs between the Cys-145 and His-41 (Fig. 1 c). As the distance to the His-41 N_e_ is substantially shorter (1.5 Å) than the one to the Cys-145 sulfur (2.3 Å), the DTT forms a sufenamide with His-41 which is a critical amino acid for the catalytic mechanism of the 3CLpro.

**Figure 2.**
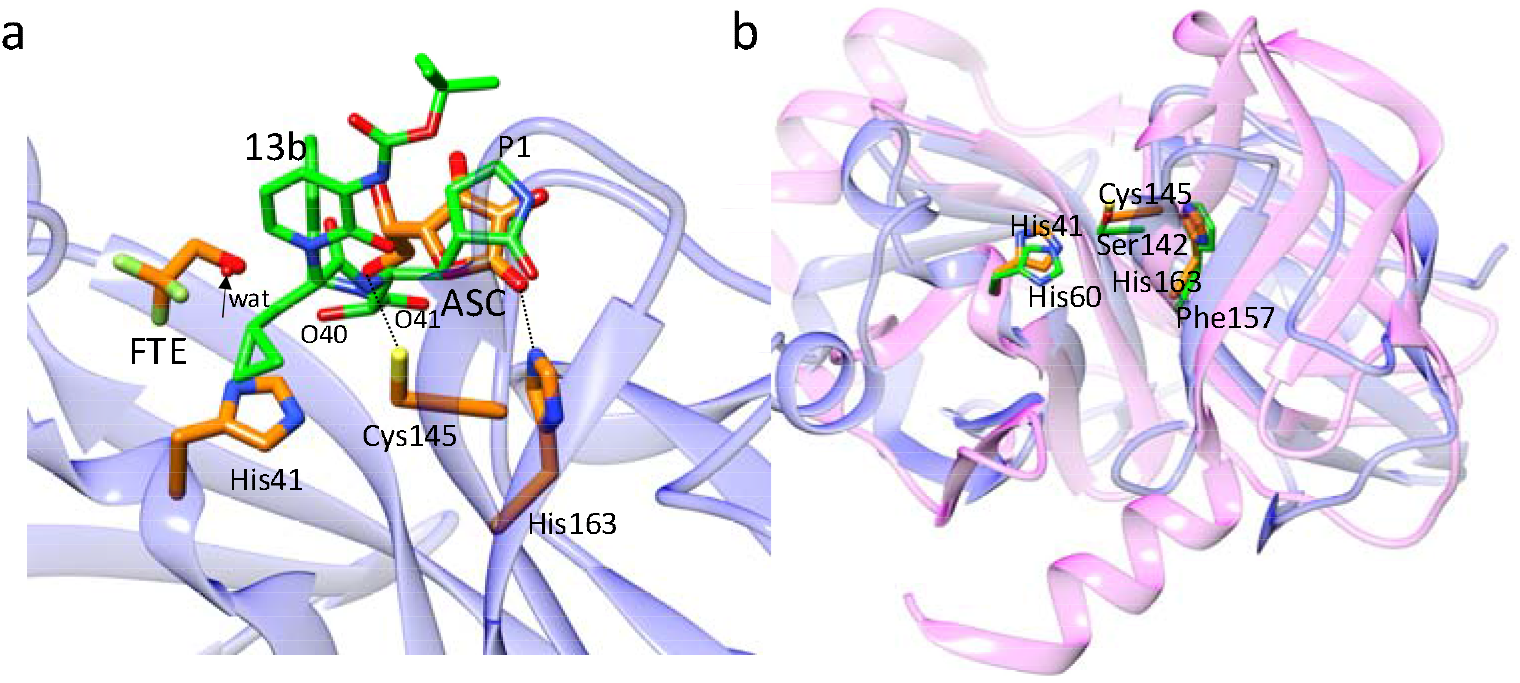
Structural comparisons. **(a)** Comparison of ligands in the active site of CoV-2 3CLpro. The position of the L-ascorbate molecule relative to compound 13b in 3CLpro as determined by Zhang et al., 2020. The positions of the C2’-oxygens is close to the two compound 13b oxygens O40 and O41 (marked), and the dihydroxyfuranone-ring of ASC coincides with the P1 moiety of compound 13b. The arrow points to a water molecule found in the structure of 3CLpro complexed with compound 13b. This position is occupied by the FTE oxygen. **(b)** Overlay of the N-terminal domain of CoV-2 3CLpro (blue ribbon, from residues 8 to 181) on the Hepatitis-C virus (HCV) serine protease (magenta ribbon). Active site residues are shown (orange for the 3CLpro, and green for the HCV protease). The active sites of both proteins are very similar. The positions of the catalytically active Cys145/Ser142 and His41/His60 are essentially identical. His163 in CoV-2 3CLpro is replaced by the structurally similar Phe157 in the HCV protease.

**Table 1.**
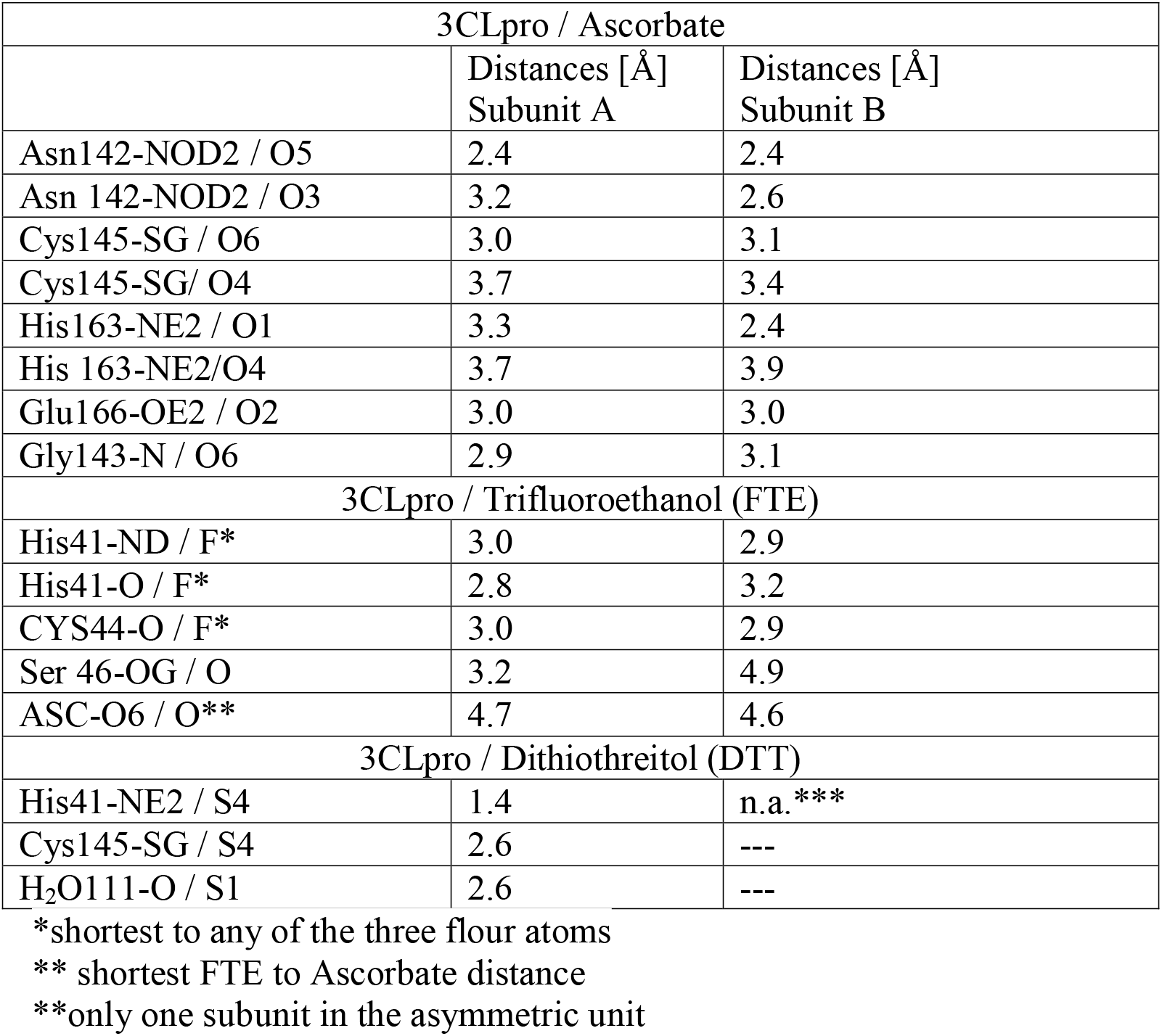
Comparison ligand interactions with conserved amino acids in the active site of subunits A and B. Designated atoms of amino acids in subunits A and B and ascorbate ligands that interact are labelled according to annotations in the refined structure (see also Scheme 1).

A search for ascorbate in the protein data bank yields 645 protein structures with only 40 structures that have a bound ascorbate. None of the 40 structures are viral proteins. To our knowledge, this is the first structure of a viral protein with ascorbate bound in the active site. The CoV-2 3CLpro has homologues in many other viruses different from coronavirus. As mentioned above, a similar (3C) protease can be found in picornaviruses (Ramajayam et al., 2011). Members of the *picornaviridae* family cause diseases such as polio (poliovirus), encephalomeningitis (encephalomyocarditis virus) and the common cold (rhinovirus) (Sharma & Gupta, 2017). Moreover, the structure of the N-terminal domain of the CoV-2 3CLpro, as well as that of its active site is very similar to the structure of the hepatitis-C virus (HCV) protease (Fig. 2), except that the HCV protease is a serine protease. However, even the HCV protease Ser-142 oxygen position is essentially identical to that of the Cys-145 sulfur in the 3CLpro. The binding of Vitamin C to the CoV-2 3CLpro might point to reasons why Vitamin C could be beneficial for the treatment of coronavirus caused diseases such as SARS and middle east respiratory syndrome (MERS) who have structurally similar 3CLpros (Hilgenfeld, 2014; Zhang, Lin, Sun, et al., 2020). Moreover, Vitamin C might also bind to proteases of other viruses, and assists in the treatment of diseases mentioned above and others such as AIDS (Harakeh et al., 1990), herpes (Furuya et al., 2008; Hoog et al., 1997), rhinovirus induced respiratory distress syndrome (Fowler Iii et al., 2017) and even rabies (Banic, 1975).

The available inhibition test (the ‘Untagged (SARS-CoV-2) Assay Kit’, BPS Biosciences) cannot be used to determine an inhibitory effect of L-ascorbate as the ascorbate quenches the fluorescence of the enzymatically produced product at higher concentrations and inhibition tests are inconclusive. Since ascorbate is detected in the crystal structure, it may well be that inhibition of proteases with Vitamin C, or perhaps with one of its derivates (Mescic Macan et al., 2019) that may even bind directly to the catalytically important Cys-145, becomes an important factor in the treatment of Covid-19 cases and maybe other virus caused infections (Colunga Biancatelli et al., 2020). Further investigations are necessary.

## Supplementary Material

### Expression

The CoV-2 3CLpro sequence was synthetized (GenScript) for optimized expression in E. coli according to sequence information published previously(Zhang, Lin, Sun, et al., 2020). In short, the N-terminus of 3CLpro is fused to glutathione-S-transferase (GST). It further has a 6-His tag at the c-terminus. The N-terminal GST will be autocatalytically cleaved off after expression due to an engineered 3CLpro cleavage sequence. Although the His tag can be cleaved off by a PreScission protease, the tag did not interfere with crystallization and consequently was left on. Overexpression and protein purification protocols were modified from previous reports. E. coli were grown to 0.8 OD_600_ at 37° in terrific broth. Expression was induced by 1 mmol/L IPTG at 25° C. After 3 h of expression, the culture was induced a second time (1 mmol/L IPTG), and shaken overnight at 20° C. The yield is about 80 mg for a 6 L culture. Cells were resuspended in lysis buffer (20mM Tris Base, 150 mmol/L NaCl, pH 7.8.). After lysis of the bacterial cells, debris was centrifuged at 50,000 g for 1 hour. The lysate was let stand at room temperature for 3 h (overnight is also possible). After this, the lysate was pumped through a column containing 15 mL of Talon Cobalt resin (TAKARA). The resin was washed without using imidazole using a wash cycle consisting of low salt (20 mmol/L Tris Base, 50 mmol/L NaCl, pH 7.8), high salt (20 mmol/L Tris Base, 1 mol/L NaCl, pH 7.8) and low salt (as above) solutions (about 20 column volumes each). After the wash cycle was completed, the column was let stand for an additional 2 h at room temperature followed by another wash cycle. The final product was eluted by 300 mmol/L imidazole. Buffer exchange was achieved at 4 ° C by either by 3 times spin-concentration and re-dilution with 20 mmol/L Tris base, 150 mmol/L NaCl, 25 mmol/L Na-ascorbate, or by dialyzing immediately in 20 mmol/L Tris base, 150 mmol/L NaCl, 0.1 mmol/L dithiotreitol (DTT), pH 7.8. The preparations were concentrated to a 3CLpro concentration of 20 mg/mL. Since the 6-His tag was not cleaved off, this one step purification protocol required only ~10 h from cell lysis to the pure 3CLpro product. The product is within 1.7 Da of the theoretical molecular weight as determined by mass spectroscopy.

### Crystallization

The concentrated 3CLpro containing DTT was diluted to 4 mg/mL. 100 μL of the diluted 3CLpro was mixed (1:1) in batch mode with the same amount of 25 % PEG 3350, Bis-Tris 100 mmol/L, pH 6.5. Rectangular shaped crystals with dimensions of about 200 × 30 × 30 μm^3^ were obtained. Crystals of the ascorbate containing 3CLpro were obtained by the hanging drop method by mixing 2 μL of 30 mg/mL protease with an equal amount of a reservoir solution containing 15 % PEG 3350, 5 mmol/L ascorbate, and trifluoroethanol (FTE) (4 %) as an additive. Crystals formed long thin needles after 3-day incubation at 16°C. Crystals were soaked in mother liquor containing an addition of 120 mmol/L ascorbate and 20 % glycerol as a cryoprotectant for 3 min before freezing.

### Data Collection and Structure determination

The crystals were mounted in Mitegen micro-loops (30 - 50 μm) and directly frozen in pucks suspended in liquid nitrogen for automated (robotic) data collection. The dewar with the pucks were provided to the Advanced Photon Source, Argonne National Lab, Lemont, IL, for robotic data collection at Sector-19 (Structural Biology Center, SBC, beamline 19-ID-D). Data collection was fully remote due to restriction of the COVID-19 pandemic. Dataset to 2.2 Å and 2.5 Å, respectively, were collected (0.5° rotation and 0.8 s exposure per detector readout for a total of 180°) on the 3CLpro with DTT and L-ascorbate, respectively. Data was processed with HKL3000 (Minor et al., 2006). Data statistics in shown in Tab. 1. The spacegroup of the DTT containing crystals was C2. For refinement, the 3CLpro structure with pdb access code 6Y2E (Zhang, Lin, Sun, et al., 2020) was used as initial model. Molecular replacement was not necessary as the model fits immediately. Refinement was achieved using refmac (Murshudov et al., 2011) (version 5.8.0238). The electron density of the DTT becomes apparent by a difference electron density feature in between His-41 and Cys-145 (Fig. 1 c). With L-ascorbate and FTE in the buffer, the spacegroup becomes P2_1_2_1_2_1_ with cell constants not found so far for the CoV-2 3CLpro. Molecular replacement was performed by Phaser (Oeffner et al., 2013) version 2.8.2 using pdb-entry 6XQT (Kneller et al., 2020) as a search model. After refinement of the molecular replacement solution, difference electron density of both, the L-ascorbate and the FTE becomes apparent in the active sites of both subunits (Fig. 1 a). The positions and orientations of the L-ascorbates and the FTE molecules were determined with the help of Polder maps (Fig. 1a and b) (Liebschner et al., 2017) calculated using Phenix (Adams et al., 2010) v1.19.2-4158. After conventional refinement in Phenix, grouped occupancy refinement resulted in sub-stochiometric concentrations with occupancy values of the L-ascorbates varying from ~60 % in subunit A to ~70 % in subunit B.

---

**Table S1.** Data collection and refinement statistics.

|  |  |  |
| --- | --- | --- |
|  | SARS CoV-2 3CLpro |  |
| Beamline | APS 19-ID-D |  |
| Temperature | 110 K |  |
| Ligand | L-ascorbate (Vitamin C) | Dithiothreitol (DTT) |
| Resolution (Å) | 2.3 | 2.3 |
| Space group | P2<sub>1</sub>2<sub>1</sub>2<sub>1</sub> | C2 |
| Unit-cell parameters (Å, °) | a = 65.4 Å b = 67.5 Å c = 157.5 Å<br>$\alpha=90^\circ$ $\beta=90^\circ$ $\gamma=90^\circ$ | a = 113.39 Å b = 53.4 Å c = 44.7 Å<br>$\alpha=90^\circ$ $\beta=102.8^\circ$ $\gamma=90^\circ$ |
| No of unique reflections | 30845 | 9426 |
| Redundancy | 5.7 (3.9) | 3.1 (1.8) |
| Completeness (%) | 98.2 (90.0) | 70.0 |
| CC1/2 @ dmin | 45.2 | 47.6 |
| R<sub>merge</sub> (%) | 20.4 (145.6) | 7.3 (122) |
| Model Building/<br>Refinement | Phenix 1.19.2-4158<br>to 2.3 Å | refmac 5.8.0258<br>to 2.2 Å |
| R<sub>cryst</sub>/R<sub>free</sub> (%) | 21.0/24.4 | 19.1/26.3 |
| # subunit/asymmetric unit | 2 | 1 |
| # residues/subunit | 303/310* | 303 |
| additional ligand | trifluoroethanol (FTE) | n.o. |
| ligand occupancy | Vitamin C: A 0.66; B 0.67**<br>FTE: A 0.66; B: 0.68 | 0.85 |
| # water | 198 | 1184 |
| R.m.s.d. bonds (Å)/angles (°) | 0.005/1.07 | 0.007/0.949 |
| Coordinate Error (Å) | 0.25 | 0.18 |

\*in subunit B, electron density for seven additional C-terminal residues including that of two histidines of the hexa-his tag can be observed.
\*\*the difference electron density in any $F^{\text{obs}} - F^{\text{calc}}$ difference map including the Polder map is visually weaker in the active site of subunit A compared to that of subunit B.
n.o.: no additional ligand

**Figure S1.**
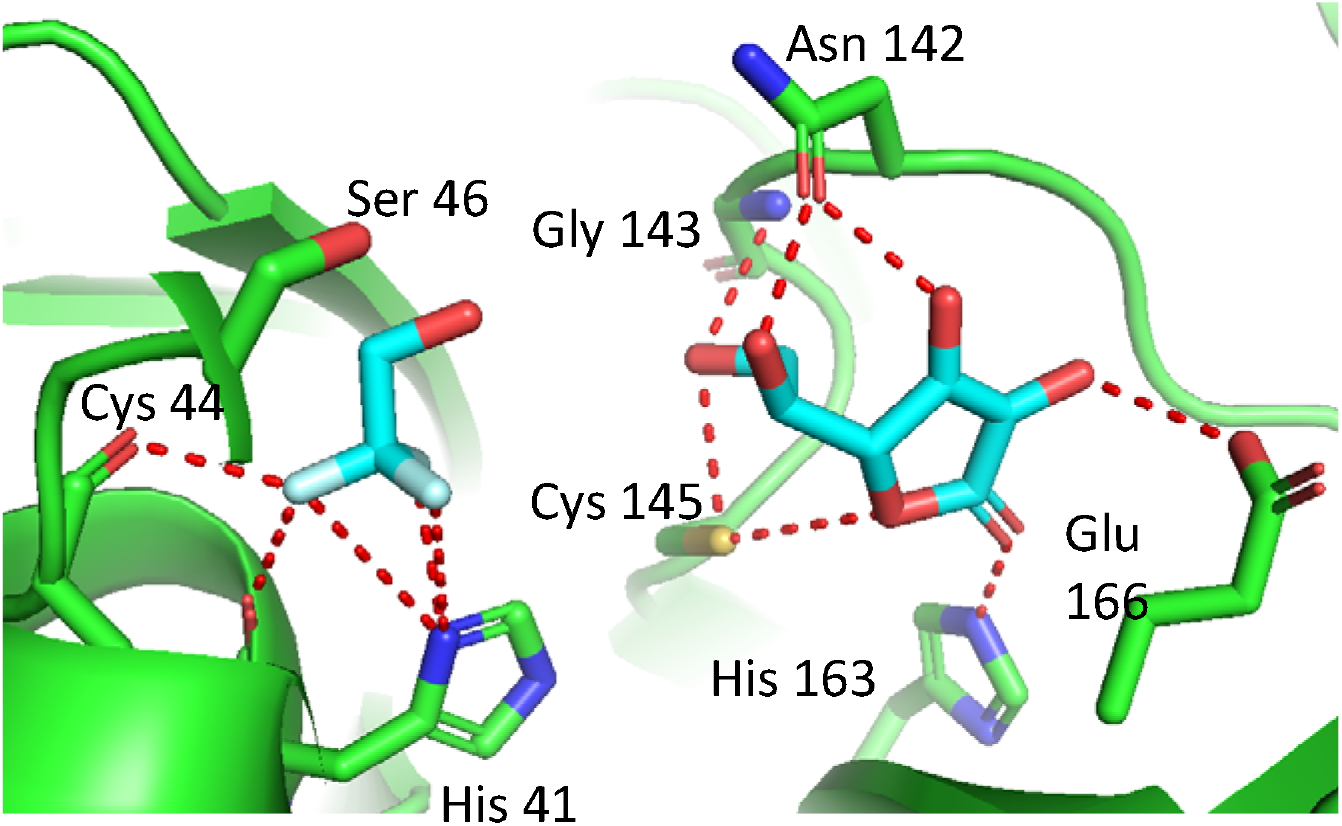
Interaction pattern of the L-ascorbate and FTE with the active site of 3CLpro subunit B. The pattern is very similar in subunit A.

